# Cortactin stabilizes actin branches by bridging activated Arp2/3 to its nucleated actin filament

**DOI:** 10.1101/2023.08.16.553490

**Authors:** Tianyang Liu, Luyan Cao, Miroslav Mladenov, Antoine Jegou, Michael Way, Carolyn A. Moores

## Abstract

Regulation of the assembly and turnover of branched actin filament networks nucleated by the Arp2/3 complex is essential during many cellular processes including cell migration and membrane trafficking. Cortactin plays a key role in stabilizing actin filament branches by interacting with the Arp2/3 complex and actin filaments via its N-terminal Acidic domain (NtA) and 6.5 central unstructured 37 amino acid repeats, respectively ^1^, but the mechanism of this is unclear. We determined the structure of cortactin-stabilized Arp2/3 actin branches using cryo-electron microscopy. We find that cortactin interacts with the new daughter filament nucleated by the Arp2/3 complex at the branch site rather than the initial mother actin filament. Cortactin preferentially binds activated Arp3 in contrast to other nucleation promoting factors (NPFs) ^2,3^. Cortactin also stabilizes the F-actin-like interface of activated Arp3 with the first actin subunit of the new filament, and its central repeats extend along successive daughter filament subunits. Cortactin binding to Arp3 is incompatible with NPF interaction and its preference for activated Arp3 explains why it is retained at the actin branch. Our data have uncovered why cortactin displaces NPFs, while at the same time promoting synergy to regulate branched actin network dynamics.

---

The actin cytoskeleton can form many different types of dynamic supramolecular arrays - from linear bundles to branched actin filament networks which underlie its functional diversity and adaptability ^4-9^. Distinct F-actin arrays are formed by the localized activities of specific actin nucleating factors, actin binding proteins and myosin motors ^4,5,10^. For example, branched actin networks are generated when a new “ daughter” filament is nucleated from the side of a pre-existing “ mother” filament by the Arp2/3 complex ^5,11,12^. Branch formation requires activation of the seven subunit Arp2/3 complex which involves a conformational rearrangement of the complex so that a short-pitch helical F-actin-like template is formed by the <u>A</u>ctin <u>r</u>elated <u>p</u>roteins Arp2 and Arp3 from which the fast growing barbed end of the daughter filament extends ^13,14^. Class 1 nucleation promoting factors (NPFs) such as WAVE and WASP activate Arp2/3 via a conserved C-terminal VCA domain, consisting of one to three **<u>V</u>**erprolin domains (also known as WASP-homology 2 domains) followed by **<u>C</u>**entral and **<u>A</u>**cidic segments ^2,3,15-21^. The VCA domain of Class 1 NPFs also stimulate nucleation by recruiting actin subunits to the activated Arp2/3 complex, from which these NPFs are subsequently released (*Extended Data Fig. 1a*) ^22-24^. The correct functioning of Arp2/3-nucleated branched actin networks depends not only on their spatial and temporal assembly but also their stability and turnover ^25-29^. The actin binding protein cortactin, a Class 2 NPF, plays a major role in stabilizing actin branches by interacting with the Arp2/3 complex and F-actin (**Fig. 1a**) ^30-33^. Furthermore, although cortactin by itself can weakly activate the Arp2/3 complex, it synergizes with Class 1 NPFs to further stimulate efficient Arp2/3-mediated actin branch formation ^22,23,34^. Given its central role in stabilizing branched actin networks, cortactin is important in many cellular processes such as epithelial integrity and intracellular trafficking as well as a range of pathologies including bacterial infection and cancer metastasis ^1,35-37^. However, despite its functional importance, the precise mode of action of cortactin and its mechanism of synergy with Class 1 NPFs remains unknown.

**Figure 1.**
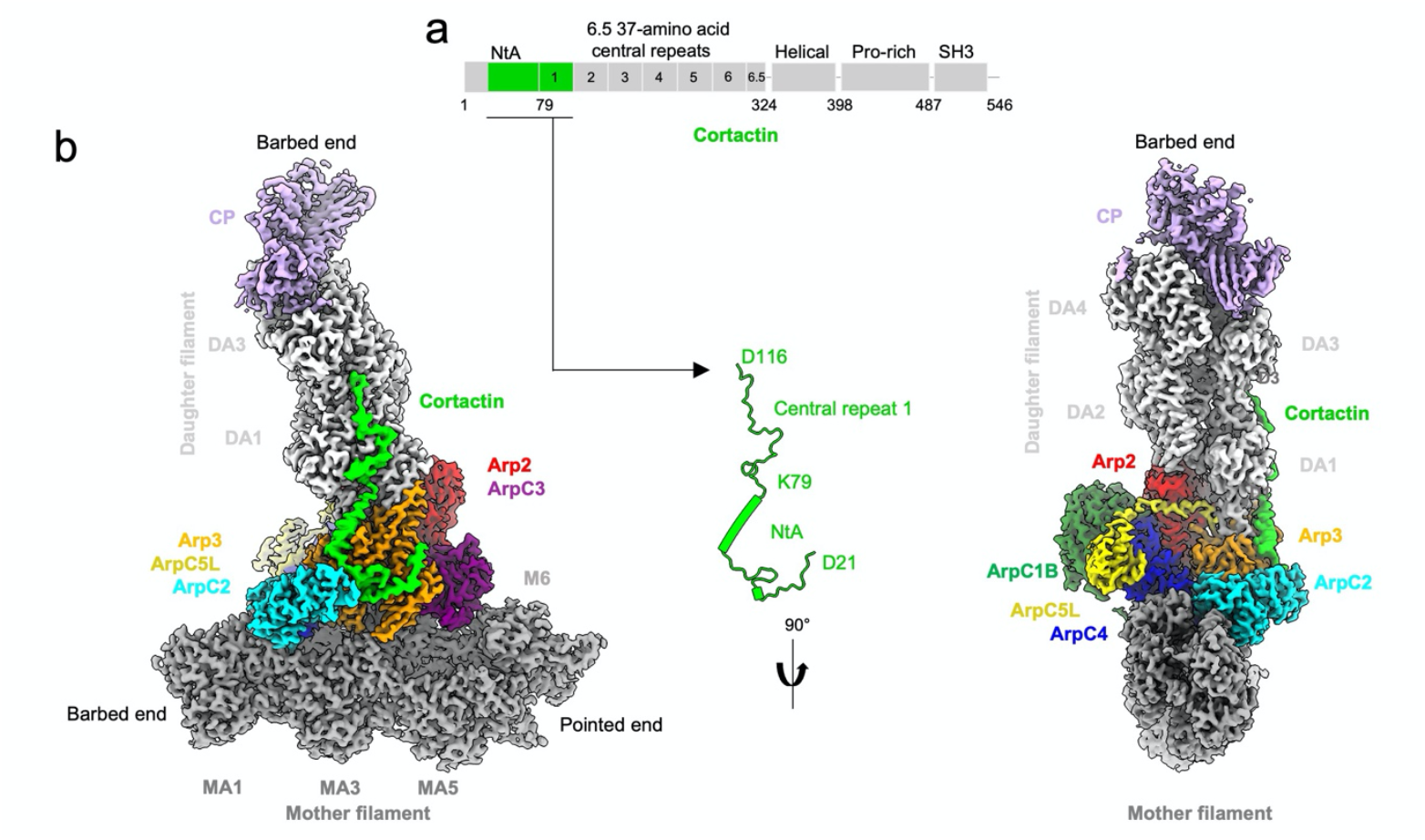
Cortactin binds the daughter filament at Arp2/3-mediated actin branches. **a)** Cortactin domain organisation. **b**) Overview of the composite cortactin-stabilized Arp2/3 actin branch cryo-EM reconstruction, assembled from four local refined reconstructions as shown in Extended Data Fig. 2 and 3. Density of individual proteins in the complex are coloured according to the labels, while mother and daughter filament subunits are coloured dark and light grey and labelled MA1, MA3, MA5 and MA6 and DA1 – DA4 respectively. The free barbed and pointed ends of mother and daughter filaments are also labelled. Central inset shows the cortactin model calculated from the cryo-EM reconstruction, with the visualized regions mapped on to the cortactin schematic in a) as indicated.

We determined the structure of *in vitro* reconstituted cortactin-stabilized Arp2/3 actin branches using cryo-electron microscopy (cryo-EM) and single particle reconstruction at ∼3.3 Å resolution (**Fig. 1 b**, *Extended Data Fig. 1, 2, Extended Data Table 1, Movie 1*). To maximize the number of branches in our sample and therefore the possibility of visualizing the previously elusive binding site of cortactin, we used the most active isoform of human Arp2/3 (Arp2/3-C1B-C5L ^33^), and also included capping protein in our sample to limit daughter filament growth ^38-41^. The resulting branch structure allowed us to visualize cortactin (*Extended Data Fig. 2, 3*) and showed that unexpectedly, cortactin connects the activated Arp2/3 complex and the daughter filament, in contrast to previous proposals that cortactin binds to the mother filament (**Fig. 1 b**) ^23,34^. In the presence of cortactin, the overall conformation of the activated Arp2/3 complex at the junction of mother and daughter filaments is similar to previous cryo-EM structures ^13,42,43^ *(Extended Data Fig. 4a, b, Movie 1*). The daughter filament consists of 4 subunits (DA1 - DA4), each with ADP bound, and its barbed end is terminated by capping protein ^38-41^. The cortactin density that extends along this short daughter filament corresponds to the first cortactin repeat (**Fig. 1b**). No density corresponding to cortactin is observed on the ADP-bound mother filament (consisting of MA1 – MA6 in our image processing scheme, *Extended Data Fig. 2*). Both mother and daughter filaments adopt canonical ADP-F-actin structures ^44,45^ and Arp2 and Arp3 are also bound to ADP (*Extended Data Fig. 4b-d*). Overall, our structure shows that rather than modifying the filaments at actin branches, cortactin branch stabilization is mediated by the protein-protein contacts that cortactin forms with the activated Arp2/3 complex and the daughter filament ^30-32^.

**Figure 2.**
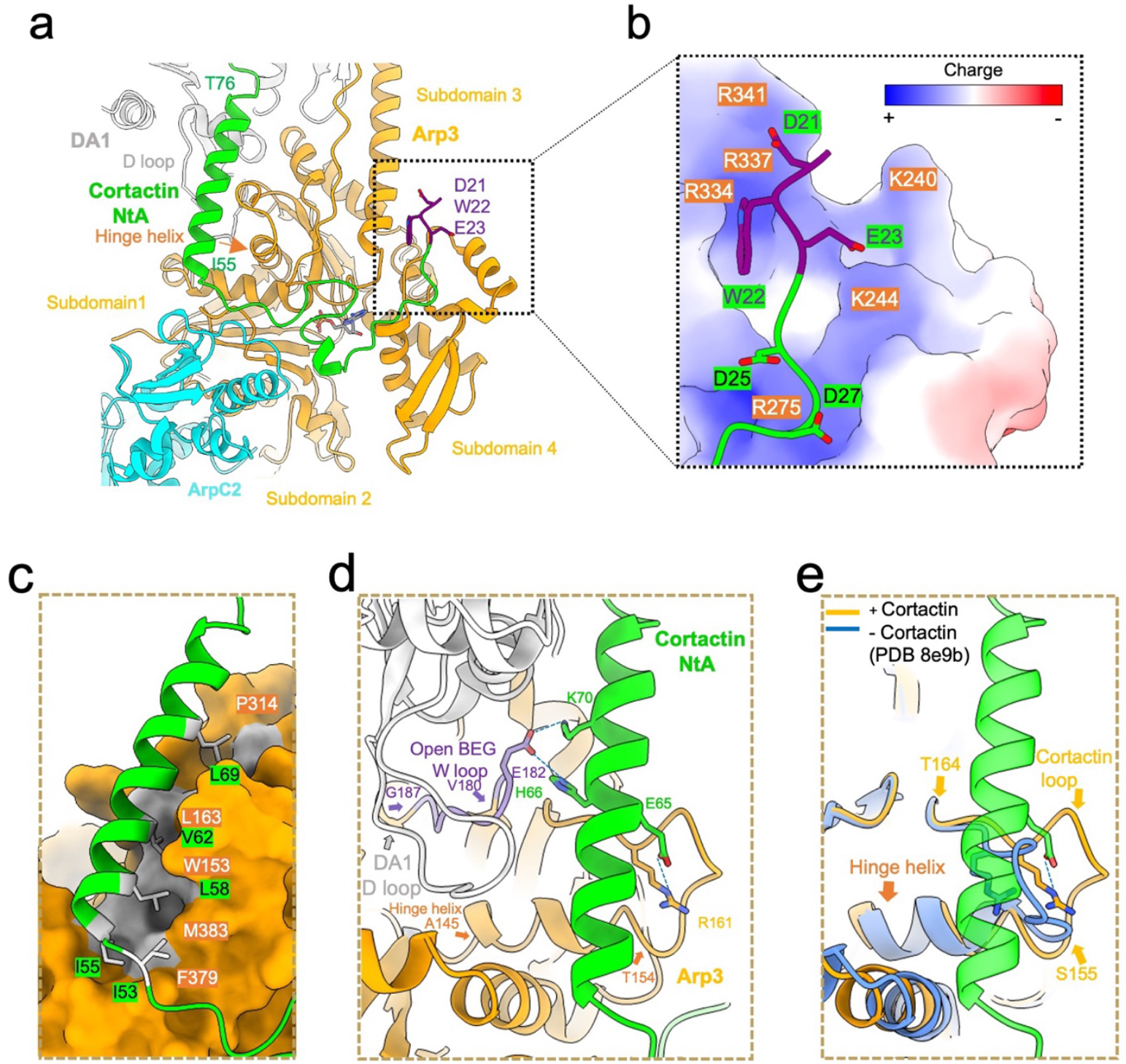
Cortactin NtA binds to activated Arp3. **a)** Overview of cortactin NtA (lime and purple) and its interactions with Arp3 (orange) and ArpC2 (cyan). DWE motif residues are shown in purple stick representation, while the rest of cortactin NtA is depicted in ribbon representation. Cortactin residues D21 - N54 meander across the Arp3 surface while residues I55 - T76 form an amphipathic *α*-helix. Subdomains in Arp3 are labelled and the Arp3 hinge helix at the junction of subdomain 1 and 3 is indicated with an orange arrow. DA1 is the first subunit of the daughter filament which via the D-loop in its subdomain 2, forms longitudinal contacts with Arp3. Detailed views of interactions are shown in *Extended Data Fig. 5a-d*. **b)** Electrostatic interaction of the negatively charged cortactin N-terminal region that inserts into a positively charged pocket of Arp3. Arp3 is depicted in surface representation, acidic regions in red, basic regions in blue with individual basic residues indicated in orange. Cortactin is depicted in ribbon model, D21-W22-E23 motif in purple stick representation, residues 24 - 29 in lime with acidic residues shown in stick representation. **c)** Hydrophobic interaction of the cortactin NtA *α*-helix (green ribbon model) with a hydrophobic groove on the surface of Arp3 (orange space-filling representation, with hydrophobic regions in grey). Residues on the interaction surface of cortactin and Arp3 are labelled in lime and orange respectively. **d)** The cortactin NtA *α*-helix stabilizes the activated Arp3 W-loop (residue 180 – 187, purple) conformation to enable DA1 actin subunit binding via its D-loop in the open barbed end groove (BEG) of Arp3. Residues forming salt bridges are shown in stick representation. The distances between interacting residues are shown in Extended Data Fig. 5a-d. **e)** The cortactin NtA *α*-helix (lime) stabilizes and interacts with a specific conformation of the Arp3 loop (residues 155 - 164), which we term the cortactin loop (in orange). This loop conformation is distinct in the presence of cortactin and is different in activated Arp2/3 in the absence of cortactin (blue ribbon).

**Figure 3.**
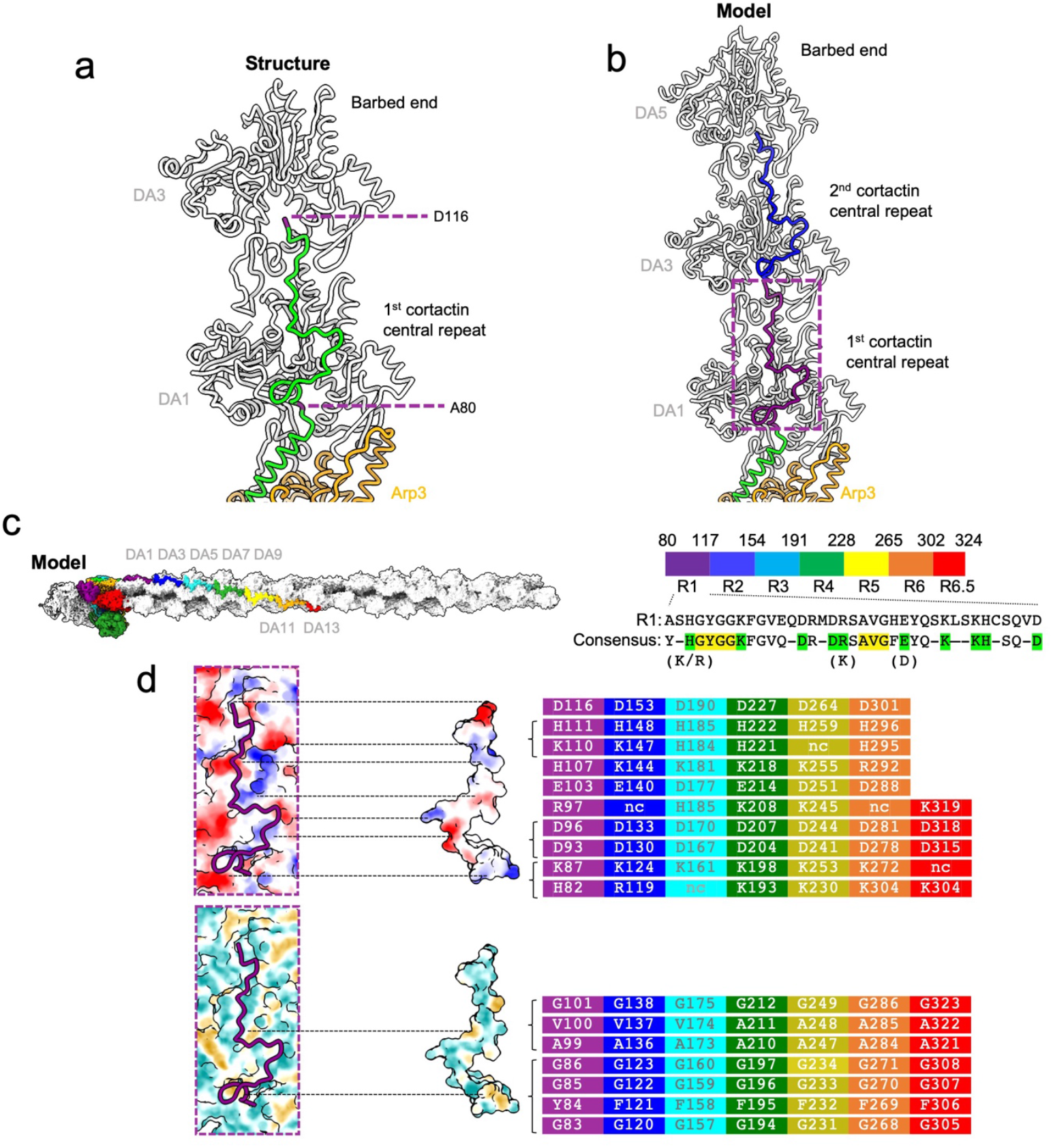
Cortactin repeats bind the daughter filament. **a)** The first cortactin repeat (in green) binds longitudinally along the daughter filament and bridges between DA1 and DA3 actin subunits (in grey). The first and last residues of the first cortactin repeat are labelled, coloured in purple and their positions on DA1 and DA3 subunits respectively are circled in purple. See also *Movie 2*. **b)** Cortactin repeats are predicted to bind equivalent subunit positions along the same strand of the daughter filament. The first repeat that bridges between DA1 and DA3 subunits is shown in purple (as in a)) and the second repeat that bridges between DA3 and DA5 is shown in blue. **c)** Left: The 6.5 cortactin repeats are predicted to bind longitudinally along 7 daughter actin filament subunits. The modelled cortactin central repeats are coloured from N-to C-terminus in purple, blue, cyan, green, yellow, orange and red. Right: Amino acid sequence of 1^st^ repeat and the consensus sequence of the 6.5 central repeats. Amino acid residues present in more than five of the 6.5 repeats are included in the consensus sequence. Charged residues (K/H/R and D/E) are grouped together in the analysis. Conserved residues in green form potential electrostatic interactions with actin subunits. Conserved residues in yellow form potential hydrophobic interactions with actin subunits as indicated in d). **d)** Top: Binding surface of DA1-actin and cortactin first central repeat coloured by electrostatic potential and shown in open book representation. Blue, positively charged; Red, negatively charged. Conserved charged residues in cortactin are shown and coloured according to the individual central repeat they are in as in Fig. 3c. Dotted lines indicate interaction regions in the assembly. Bottom: Binding surface of DA1-actin and cortactin first central repeat coloured by hydrophobicity and shown in open book representation. Yellow = hydrophobic; Cyan = hydrophilic. Conserved hydrophobic residues are shown and coloured according to the individual central repeat they are in as in **Fig. 3c**; nc = not conserved. Dotted lines indicate interaction regions in the assembly.

**Figure 4.**
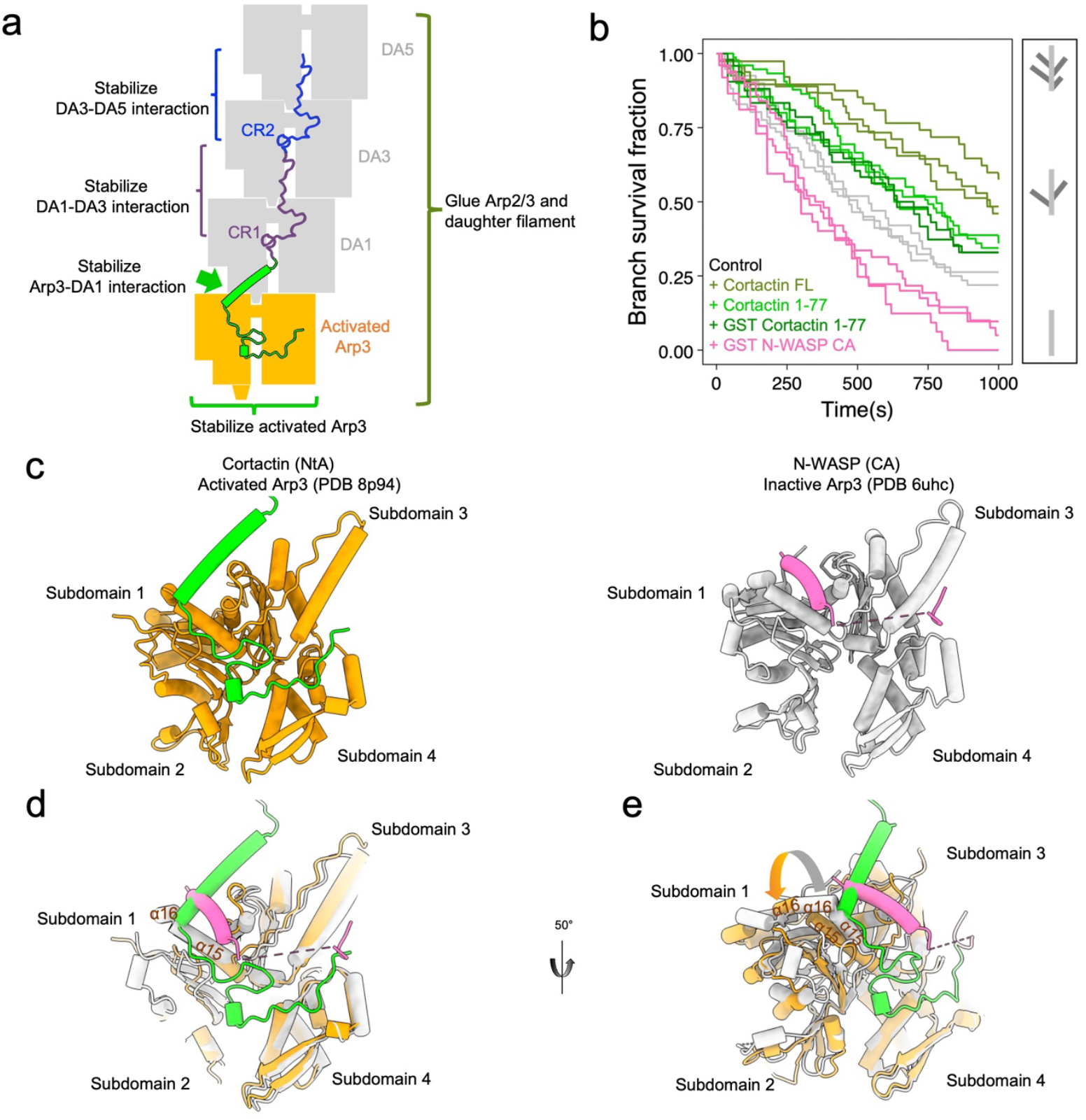
The binding site of NtA cortactin on Arp3 explains its synergy with VCA domains. **a)** Schematic showing how cortactin stabilizes the actin branch junction. **b)** The survival fraction of Arp2/3 mediated branches over time. The dissociation of Arp2/3 mediated branches was observed and quantified in the presence of 0.1 µM full length cortactin (olive green), or in the presence of 0.1 µM cortactin NtA (dark green for GST-NtA and light green for untagged NtA), or in the presence of 0.1 µM GST-N-WASP-CA (pink) in addition to 0.3 µM G-actin. The results of the control experiments with only 0.3 µM G-actin are shown in grey. Data for each curve were obtained from independent experiments. Schematic on right indicates actin branch survival status in the assay (mother filament dark grey, daughter filaments light grey). **c**) Binding sites of cortactin NtA (left, green) and N-WASP-CA (right, pink) on active Arp3 (orange) or inactive Arp3 (grey). Arp3 subdomains are numbered. **d)** Overlapping binding sites of cortactin NtA and VCA on Arp3 indicate how these proteins would compete for Arp3 binding. Active/inactive Arp3 structures are superposed by alignment of subdomains 3 and 4. Only a subset of Arp3 structural features are shown for clarity. **e)** Rotated view of overlaid active (orange) and inactive (grey) Arp3 structures with cortactin NtA and CA domain bound, as in (d). Conformational differences of Arp3 *α*-helices at the cortactin NtA and CA binding sites in the active/inactive Arp3 are indicated by arrows and explain the sensitivities of these binding partners to Arp3 activation state.

Cortactin NtA domain residues 21 - 79 form electrostatic and hydrophobic interactions with all four Arp3 subdomains, as well as contacting ArpC2 (**Fig. 2a**, *Extended Data Fig. 5a-d*). The cortactin DWE motif (residues 21 – 23) that is essential for the interaction with Arp2/3 ^31^ inserts into a positively charged pocket of Arp3, while residues 24 – 54 adopt a meandering trajectory across the Arp3 surface (**Fig. 2 a, b**, *Extended Data Fig. 5*). At residue 55, the NtA domain turns ∼90° on the surface of Arp3 and forms an amphipathic *α*-helix (residues I55 – T76) which binds in a hydrophobic cleft on Arp3 and points towards the daughter filament (**Fig. 2a, c**). This cortactin helix binds adjacent to the Arp3 hinge helix (residues 145 – 154) which is key in mediating inactive-active Arp2/3 complex structural transitions ^13,14,43^.

The cortactin NtA *α*-helix stabilizes the Arp3 W-loop (residues 180 – 187) in a conformation that has been previously observed only in activated Arp2/3 and is distinct from that seen in the inactivated Arp2/3 complex ^3,13,46,47^. As a result, the structural groove at the barbed end of Arp3 is open and promotes the interaction with DA1 of the new daughter filament (**Fig. 2d**). The contacts formed between activated Arp3 and DA1 mimic the longitudinal contacts along F-actin ^13,14^ and involve insertion of subdomain 2 of DA1 - specifically its so-called D-loop - in the barbed-end of Arp3 (**Fig. 2d**). Further, the loop within Arp3 (residues 155 - 164) that follows the hinge helix - which we now term the cortactin loop - makes contacts with the cortactin NtA *α*-helix via a distinct conformation compared to branch structures in the absence of cortactin (**Fig. 2e**). The structure is consistent with a model in which the interaction of the cortactin NtA domain with activated Arp2/3 complex also stabilizes the interface of Arp3 with DA1 of the daughter filament.

The first cortactin central repeat (residues 80 - 116) extends ∼5.5 nm from the C-terminus of the NtA *α*-helix longitudinally on the daughter filament to connect the two successive subunits DA1 and DA3 (**Fig. 3a**). D116, the final residue of the first cortactin central repeat, is positioned on DA3 in an equivalent position to the first repeat residue (A80) on DA1 (**Fig. 3a**). Based on the cortactin F-actin interaction in our structure, we modelled how the cortactin central repeats would interact with a longer daughter filament. Our model shows that the second repeat would bind along the daughter filament in the same way as the first repeat to connect DA3 and DA5 (**Fig. 3b**). Furthermore, it predicts that the cortactin repeats, including the C-terminal half repeat, would extend to the barbed end of DA13, a half-turn of the F-actin helix (**Fig. 3c**, *Movie 2*). The conservation of interacting residues within the cortactin repeats is also consistent with the repeating pattern of interactions with the hydrophobic and hydrophilic regions of the F-actin surface (**Fig. 3d**). Given the conserved amino acid distribution between all 6.5 repeats and the observed binding pattern of the first central repeat, it is likely that cortactin repeats act together to maximize branch stability. Our model shows how lysine acetylation in cortactin central repeats would reduce cortactin F-actin binding activity and thereby impede cell motility ^48^. The observation that cortactin binds exclusively along the daughter filament now also explains why cortactin stabilizes linear actin filaments nucleated by SPIN90-Arp2/3 complexes in the absence of a mother filament ^49^.

Our data show that the daughter filament is stabilized directly by the cortactin repeats binding along intra-strand subunits of the daughter filament (**Fig. 4a**). In addition, they also reveal that the NtA indirectly stabilizes the branch by forming extensive interactions with activated Arp3 to promote DA1 D-loop insertion (**Fig. 2a**, *Extended Data Fig. 5*). The interactions of cortactin NtA and Arp3 are specific to its activated conformation as computational docking of our NtA structure onto the inactive Arp3 conformation generates structural clashes (**Fig. 2a**, *Extended Data Fig. 5e*). To assess the idea that NtA alone can stabilize actin branches because of its preference for activated Arp3, we tested the ability of cortactin NtA to maintain branches in an *in vitro* debranching assay (**Fig. 4b**, *Extended Data Fig.6*). NtA does provide protection from debranching compared to the actin-only control. However, it was less effective than the full-length cortactin, consistent with the notion that central repeat binding maximizes branch stabilization.

Our observation that cortactin stabilizes activated Arp3 contrasts with VCA-containing class 1 NPFs and has implications for the coordinated regulation of actin branches. Our structure shows how binding of cortactin NtA to Arp3 would sterically block VCA binding sites - particularly of its C-helix - to compete for Arp3 binding (**Fig. 4c, d**). VCA binding to Arp3 has only been observed on the inactive complex ^2,3^, it does not readily associate with activated Arp3 in MD simulations ^50^, and VCA binding would clash with the D-loops of incoming DA1 and DA2 ^13^. Consistent with a preference for inactivated Arp2/3, N-WASP-CA promotes branch destabilization in our debranching assay ^49^ (**Fig. 4b**, *Extended Data Fig*.*6*). The overlapping binding sites of cortactin NtA helix and VCA helix are centred at the junction of Arp3 subdomains 1 and 3; the relative position of these subdomains alters on Arp2/3 activation, and each protein is sensitive to these changes (**Fig. 4d**). Further, the preferential binding of VCA and cortactin to inactive and activated Arp3 respectively provides a mechanistic basis for displacement models of the synergistic activation of Arp2/3 by VCA and cortactin. VCA release from nascent actin branches is a necessary and rate-limiting step for branch formation and is accelerated by cortactin^22-24^. Our structure shows that cortactin displaces the NPF CA domain, both by competition and because the activated Arp3 conformation favours NtA binding. The previously reported synergy of class 1 NPF VCA domains and cortactin at Arp2/3 branches therefore arises from VCA binding to and activating Arp2/3 followed by NtA accelerating VCA release and stabilizing the Arp2/3 activated state ^22,23,34^. Further, the observation that cortactin alone is only a weak activator of Arp2/3 nucleation has been puzzling and was thought to be because of its inability to recruit actin monomers to the nascent branch, unlike Class 1 NPFs ^32,51^. Our structure now shows that this weak stimulation of Arp2/3 nucleation is also because of the preference of the cortactin NtA for activated Arp3.

Our data reveal exactly how cortactin supports Arp2/3-mediated actin branches by binding and stabilizing the active conformation of Arp3 and the longitudinal subunit interactions along the daughter filament. It is also striking that our structure-based model reveals that the interaction mode of the 6.5 cortactin central repeats correspond precisely to a half-turn of the F-actin helix. In contrast, the haemopoietic specific cortactin paralogue, HS1, only has 3.5 repeats and would be predicted only to interact with DA1, DA3, DA5 and DA7, consistent with its lower affinity for F-actin ^52^. This indicates how the regulated expression of cortactin and its relatives in different tissues could tune the local dynamics of branched actin networks. Within the Arp2/3 complex, Arp3 not only forms the structural template for the nucleated daughter filament, but its conformation favours binding partners such as cortactin, and may also communicate to other cytoskeleton regulators such as the de-branching factor coronin that the complex is activated ^33^. Since actin branch turnover is critical for normal functioning of the actin cytoskeleton, our visualization of cortactin has important implications for how it protects against de-branching by coronin, whether via competition for Arp2/3 binding, protection of the daughter filament junction or both ^28,33,53-55^. This in turn could determine whether Arp2/3 complexes remain bound to mother filaments following de-branching and are thus available for further rounds of nucleation. Our discovery of an *α*-helix in the cortactin NtA and characterisation of its binding site at the junction of Arp3 subdomains 1 and 3 highlights the equivalence of this binding site to the binding cleft on actin where a large number of actin binding proteins interact and which also mediates longitudinal contacts in F-actin ^44,45,56^. This emphasizes the conserved nature of the conformational changes that both Arp3 and actin undergo during actin nucleation and polymerization, and the importance of this hotspot in both proteins for binding regulators.

## Acknowledgements

This project has received funding from the European Research Council (ERC) under the European Union’s Horizon 2020 research and innovation programme (grant agreement No 810207 to M.W. and C.A.M.). L.C. was supported by the European Union’s Horizon 2020 Marie Sklodowka-Curie individual fellowship program (H2020-MSCA-IF-101028239 – MolecularArp). A.J. was supported by the ERC (grant StG-679116). MW is supported by the Francis Crick Institute, which receives its core funding from Cancer Research UK (CC2096), the UK Medical Research Council (CC2096), and the Wellcome Trust (CC2096). Cryo-EM data collected at the Institute of Structural and Molecular Biology (ISMB), Birkbeck was on equipment funded by the Wellcome Trust, U.K. (202679/Z/16/Z and 206166/Z/17/Z). We thank N. Lukoyanova and S. Chen for electron microscope support, D. Houldershaw for computing support at Birkbeck.

## METHODS

### Protein purification

Full length mouse cortactin (1 - 546, UniProt Q60598), human SPIN90 C-terminus (267 - 715, UniProt Q9NZQ3-3), GST-tagged human N-WASP-VCA and N-WASP-CA (392 - 505 and 453 - 505 respectively, UniProt O00401) were purified following the protocol described in Cao *et al* ^49^. Cortactin NtA (1 - 77) was purified following the same method as full-length cortactin. Human Arp2/3 complex containing ArpC1B/C5L isoforms (UniProt P61160, P61158, O15143, O15144, O15145, P59998, Q9BPX5) was purified following the protocol described by Baldauf *et al* ^57^.

Mouse capping protein α1β2 (UniProt P47753 and P47757-2) were co-expressed in BL21 Star^™^ DE3 cells using a pRSFDuet-1 plasmid with N-terminal 6*His tag fused to the α1 subunit. Cells were grown at 37°C and protein expression was induced with 20 μM IPTG when OD reached 1.1. After adding IPTG, the cells were grown overnight at 16°C. The next day, cells were harvested by centrifugation at 5,000 g for 15 mins, resuspended and lysed using a high-pressure homogeniser (Avesti Emusiflex C3) in lysis buffer (50 mM Tris pH 8.0, 138 mM NaCl, 2.7 mM KCl with EDTA-free protease inhibitor (Roche)). The cell lysate was centrifuged at 49,500 × g for 30 mins to remove cell debris. The supernatant was transferred to a column with Ni-NTA Resin (Merck) and incubated for 1 hour at 4°C. The column was washed with lysis buffer and His-tagged capping protein dimer was eluted from the column in elution buffer (50 mM Tris pH 8.0, 138 mM NaCl, 2.7 mM KCl with 250mM Imidazole). The eluted proteins were concentrated to 0.5ml using Amicon Ultra-4 ml Centrifugal Filters (Millipore) and loaded onto a gel filtration column (Superdex 200 Increase 10/300 GL, GE Healthcare) on an ÅKTA system (GE Healthcare). The peak fractions containing capping protein were collected and buffer exchanged into a low-salt buffer (10 mM Tris pH 7.5,10 mM KCl and 1mM DTT). Finally, the proteins were loaded onto a 1ml HiTrap Q HP column (GE Healthcare). Capping protein heterodimers were separated from other minor protein contaminants by linear gradient elution. The linear gradient was generated by combining high-salt buffer (10 mM Tris pH 7.5, 400 mM KCl and 1mM DTT) with low salt buffer (10 mM Tris pH 7.5,10 mM KCl and 1mM DTT). β/γ non-muscle actin from purified porcine brain was purchased from Hypermol (Cat 8401-01) and reconstituted with 200 µl ultrapure water to obtain a 1 mg/ml solution in a buffer with 2 mM Tris-HCl pH 8.2, 2.0 mM ATP, 0.5 mM DTT, 0.1 mM CaCl2, 1mM NaN3 and 0.2% disaccharides.

### Cryo-EM sample preparation

Branch reconstitution conditions were adapted from those used to reconstitute *Schizosaccharomyces pombe* Arp2/3 complex-bound actin filaments ^14^. Protein concentrations were optimized to enhance short actin branch formation and minimize the preferred orientation problem caused by the “ Y” -shape of actin branches on the cryo-EM grid: 1) actin concentration was kept low to prevent spontaneous nucleation and limit filament growth; 2) a high concentration of capping protein was added to limit daughter filament growth. First, 1.7 μM Arp2/3, 1.7μM VCA, 16.1 μM SPIN90, 0.8 μM actin and 3.2 μM capping protein were mixed in 14.9 μl buffer containing 20 mM HEPES pH 7.5, 50 mM KCl, 1 mM EGTA, 1 mM MgCl_2_, 0.2 mM ATP and 1 mM DTT and incubated at room temperature for 20 mins. Then, 4.5 ul of 23.8 μM actin was added in 9 separate additions that were together incubated at room temperature for 20 mins. 1.2 μl of 80 μM capping protein was added in 2 separate additions with the 3^rd^ and 7^th^ addition of actin. After the final addition of actin, 1.7 μM cortactin was added followed by another 20 min incubation. Finally, 10 μM phalloidin (Invitrogen™) was added to stabilize the actin branches.

Following incubation, 4 μl of the final reconstitution mix was applied to a glow-discharged C-flat 1.2/1.3 grid. The grid was plunge frozen using EM GP2 Automatic Plunge Freezer (Leica) with the following settings: sensor blotting, back blotting, additional movement of 0.3 mm, blotting time of 5 s, humidity of 98%, and temperature of 22 °C.

### Cryo-EM data acquisition

Cryo-EM data (12,073 movies) were collected on a Titan Krios microscope (Thermo Fisher Scientific) operated at an accelerating voltage of 300 kV with a nominal magnification of 81K and pixel size of 1.067 Å. The data were collected with a K3 detector operating in super-resolution mode (bin2) with a BioQuantum energy filter (Gatan). 50 frames for each micrograph were collected using EPU software with 14.8 e^-^/pixel/s dose rate, 3.8 s exposure time, 49.4 e^-^/ Å^2^ total electron exposure dose and a defocus range from -0.9 to -2.4 μm.

### Cryo-EM data processing

Cryo-EM data were processed using CryoSPARC v3 ^58^. Movies were motion-corrected using Patch motion. Contrast Transfer Function (CTF) parameters were estimated using Patch CTF. 8518 micrographs with CTF fit resolution < 6.4 Å and total full-frame motion distance < 50 pixels were selected for further data processing. Blob picker with a minimum diameter of 150 Å and a maximum diameter of 200 Å was used for particle picking followed by particle extraction with a box size of 368 pixels and binning factor of 4. 2,001,580 extracted particles were subjected to multiple rounds of 2D classification to remove contaminants, carbon and non-branched portions of actin filaments. Class averages featuring various views of actin branch junction were selected as templates for template picking. 3,247, 396 template-picked particles were subjected to multiple rounds of 2D classification. The particle sets selected from the blob picker (167,590 particles) and template picker (244,162 particles) were subjected to *ab-initio* reconstruction with 2 classes respectively. After *ab-initio* reconstruction, un-binned particles from these classes were re-extracted with box size of 440 and each were subjected to Homogeneous refinement with the best branch-like *ab-initio* volume as the initial model. After homogeneous refinements and duplicate removal, the two stacks of particles were combined. The combined 179,923 particles were then subjected to a first Non-Uniform (NU)-refinement followed by Heterogeneous refinement with 3 classes to further classify particles. Class 1 volume exhibits additional density on one side of the mother filament, and these particles were discarded. The remaining 130,915 particles from class 2 and class 3 were combined and subjected to a second NU refinement. Because all 3 classes are different only on the mother filament region, we referred to the first NU-refinement reconstruction before heterogeneous refinement as the Arp2/3-daughter filament consensus map. The second NU-refinement reconstruction was referred to as the mother filament consensus map (*Extended Data Fig. 2*).

Local refinement with a mask around the mother filament on the mother filament consensus map was used to improve the mother filament density. Likewise, the Arp2/3-daughter filament consensus map was divided into three overlapping segments (the Arp2/3 complex, the daughter filament and the capping protein) and locally refined to improve the density of each segment. Before running local refinement on daughter filament and capping protein, the particles and consensus map were re-centred on DA3 using Volume Alignment Tools in cryoSPARC to improve the alignment because they are at the periphery of the consensus reconstruction. After local refinement on the daughter filament, the complete first cortactin F-actin repeat density was observed. After local refinement on capping protein, the capping protein density was well resolved. 3D classification showed that the daughter filament segment of all 3 classes had the identical feature containing only four actin subunits plus one capping protein heterodimer. A high molar ratio of capping protein to actin in our reaction mix contributes to the short daughter filament. The template picking and ab-initio reconstruction step may also bias the particle selection used in our reconstruction. Global resolution and local resolution of local refined maps are estimated in cryoSPARC (*Extended Data Fig. 3*).

### Model building

The four locally refined reconstructions were used to model Arp2/3 and cortactin NtA, daughter filament and cortactin first central repeat, capping protein and mother filament (*Extended Data Fig. 2,3*). Models of all seven Arp2/3 subunits, β actin and capping protein created with the AlphaFold Monomer v2.0 pipeline were used initially ^59,60^. They were rigidly fit into EM density using ChimeraX ^61^ followed by molecular-dynamics flexible fitting using ISOLDE^62^. Namdinator ^63^ was used to optimize bond geometry, and ISOLDE and Coot ^64^ were used at the end of the model building process to manually fix Ramachandran outliers, rotamer outliers and clashes. AlphaFold predicts the N-terminus of cortactin with low confidence except for one 6-turn α-helix, corresponding to the α-helix in our EM density. The AlphaFold-predicted α-helix (residue 55 - 76) was well fitted into the EM density with bulky side chains on one side of the α-helix facilitating its positioning. After the positioning of the NtA helix, the flanking cortactin residues (21 – 54 and 80 - 116) were manually built using Coot ^64^.

### Structural analysis and visualization

Structural figures and movies were made with ChimeraX ^61^. Rise and twist angles shown in *Extended data Fig. 4* were calculated in PyMOL Molecular Graphics System, Version 2.5.4 Schrödinger, LLC. The distance between interacting atoms in *Extended Data Fig. 5* were measured in ChimeraX.

### Dissociation of branches by cortactin and CA motifs

Microfluidics experiments were done with Poly-Dimethyl-Siloxane (PDMS, Sylgard) chambers with three inlets and one outlet based on the original protocol from Jégou *et al* ^*65*^. The microfluidic flows were monitored by a MFCS and Flow Units (Fluigent). Experiments were performed in buffer containing 5 mM Tris–HCl pH 7.0, 50 mM KCl, 1 mM MgCl_2_, 0.2 mM EGTA, 0.2 mM ATP, 10 mM DTT, 1 mM DABCO, and 0.1% BSA. The temperature was maintained at 25°C by an objective heater (Oko-lab). Actin filaments were visualized using TIRF microscopy (Nikon TiE inverted microscope, iLAS2, Gataca Systems) equipped with a 60× oil-immersion objective. Images were acquired using an Evolve EMCCD camera (Photometrics), controlled with the Metamorph software (version 7.10.4, from Molecular Devices).

Pointed end anchored mother filaments (15% labelled with Alexa-488) and their branches (15% labelled with Alexa-568) were generated in a microfluidics chamber with 20 µm height and 1600 µm width as described in Cao *et al* ^49^. During the experiment, actin branches were exposed to 0.3 µM actin as control or with an additional 0.1 µM cortactin, GST-N-WASP-VCA or their mutants. The flow rate was set as high as 16 µL/min during the measurement. The forces, ranging from 0.6 to 1 pN applied on the daughter filaments, are identical in each experiment. For each condition, the survival fraction of branches was quantified and plotted over time (**Fig. 4b**). The time points when half of the actin branches disappeared under different experimental conditions were plotted for comparison (*Extended Data Fig. 6b*).

## Data availability

The cryo-EM maps and the corresponding structural coordinates were deposited under the accession codes PDB ID: 8P94 and EMDB: 17553 (Daughter filament consensus reconstruction), 17554 (Arp2/3 complex local refined reconstruction), 17555 (daughter filament local refined reconstruction), 17556 (capping protein local refined reconstruction), 17557 (mother filament local refined map reconstruction), 17558 (mother filament consensus reconstruction).

## FIGURES

**Movie 1** – Reconstruction overview and conformational changes associated with activation of Arp2/3.

**Movie 2** – Cortactin stabilizes the daughter filament through both stabilization of activated Arp3 and stabilization of longitudinal contacts between daughter filament subunits.

