## Supplementary Figures and Table for "Cortactin stabilizes actin branches by bridging activated Arp2/3 to its nucleated actin filament"

### EXTENDED DATA

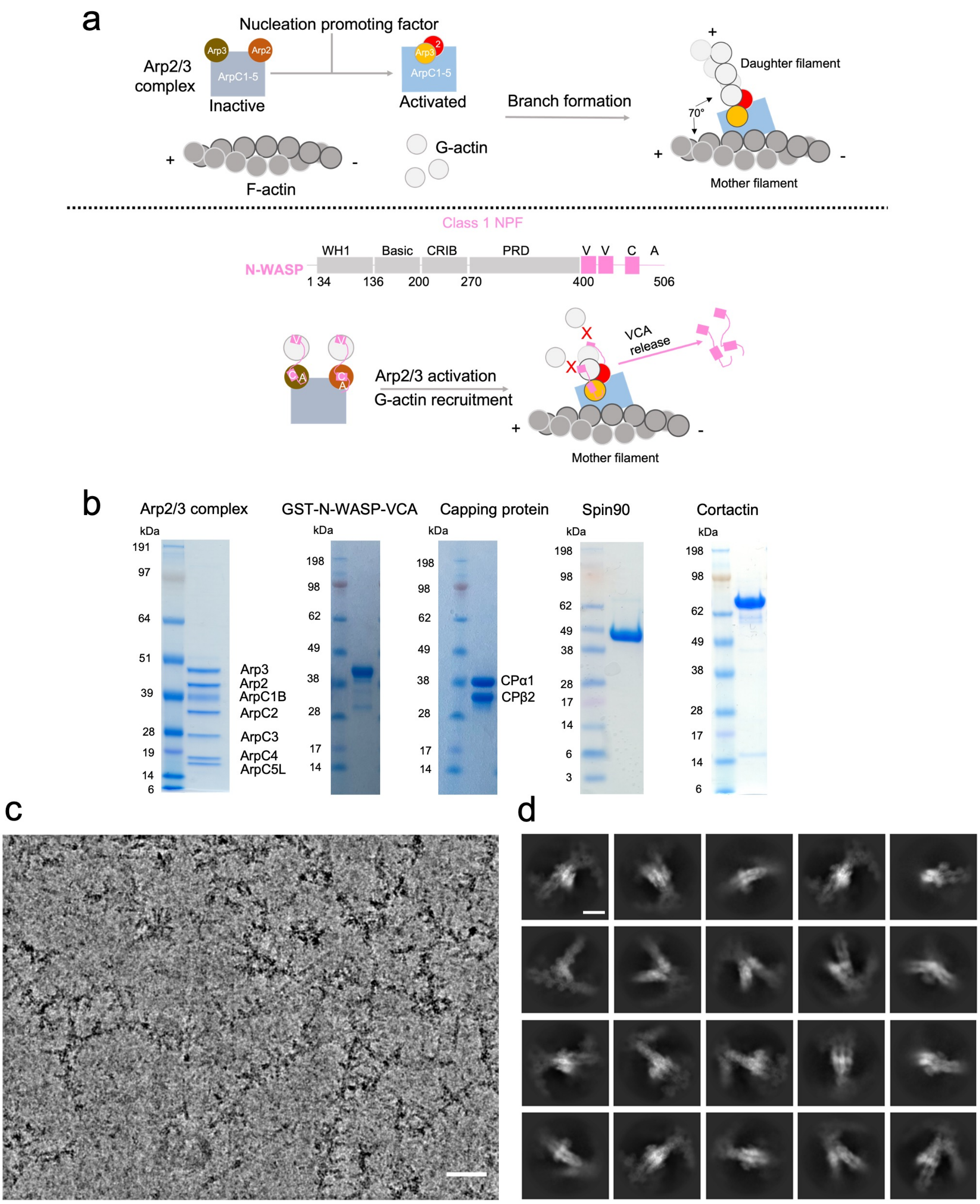


**Extended Data Fig 1. Overview of actin branch formation, purified proteins used in actin branch reconstitution and exemplar cryo-EM data.**

**a**) Schematic of Arp2/3-mediated actin branch formation and role of class 1 NPFs. When the Arp2/3 complex is activated by class 1 NPFs, its Arp2 and Arp3 subunits rearrange into a short-pitch conformation, which acts as the template for daughter filament growth. The CA domain within the VCA domain of class 1 NPF interacts with Arp2/3 and V motif binds and recruits actin monomers. VCA must be released from the nascent branch junction prior to daughter filament elongation because it blocks the binding site for further daughter filament growth. **b**) SDS-PAGE gels showing purified proteins used in cryo-EM and microfluidics reconstitution experiments. **c**) A representative cryo-EM image of cortactin stabilized Arp2/3-mediated actin branches showing "Christmas tree"-like mother filaments with multiple short daughter filaments extending from them. Scale bar = 50 nm. **d**) Representative 2D class averages of particles selected using CryoSPARC blob picker and subjected to 2D classification showing multiple 2D projection views, which are used as templates for template picking. Scale bar = 10 nm.

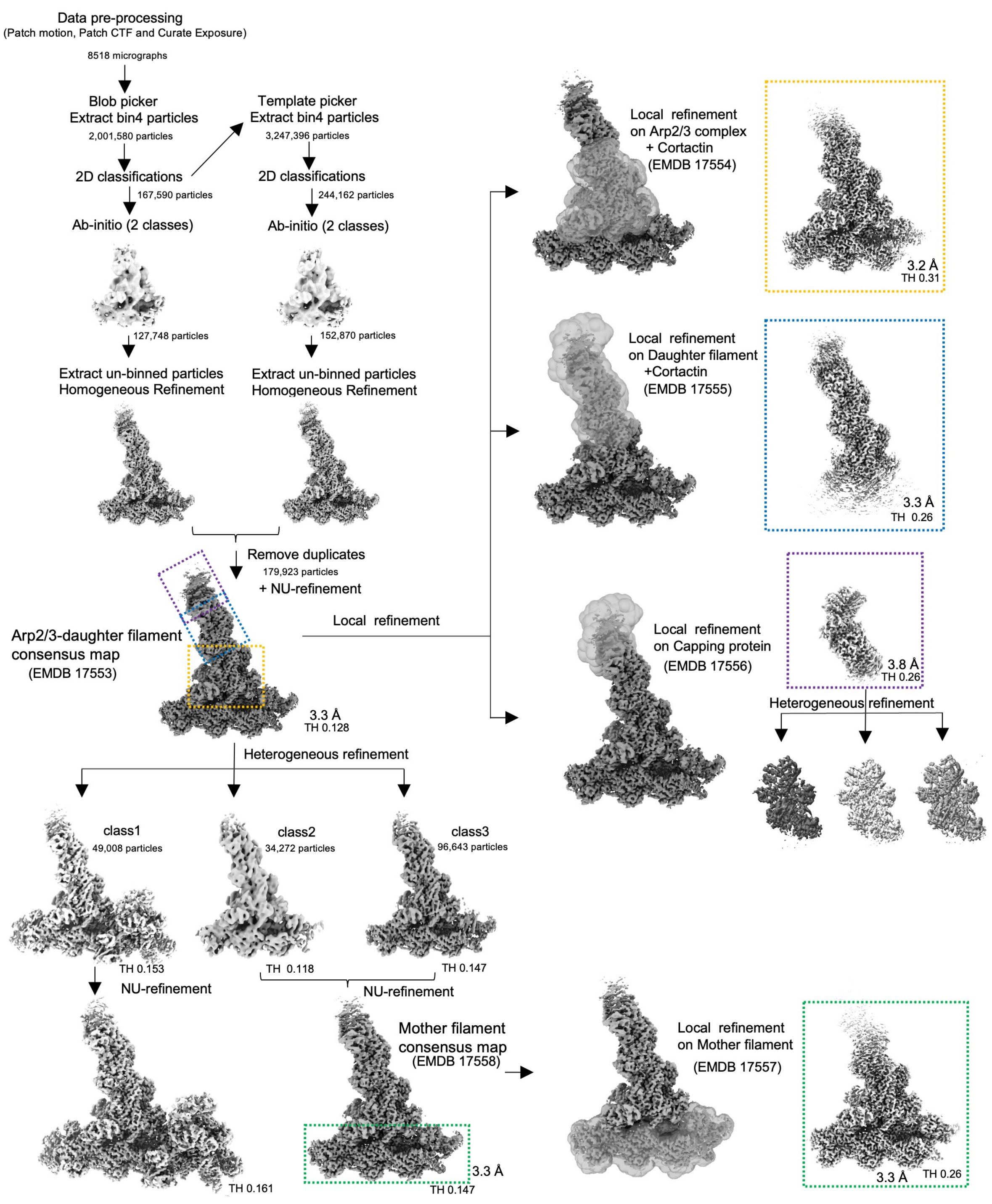


**Extended Data Fig 2. Image processing workflow for cryo-EM reconstruction.**

The workflow used to generate the overlapping locally refined reconstructions of cortactin-bound Arp2/3 complex, daughter filament, capping protein and mother filament. Thresholds (THs) and global resolutions are indicated.

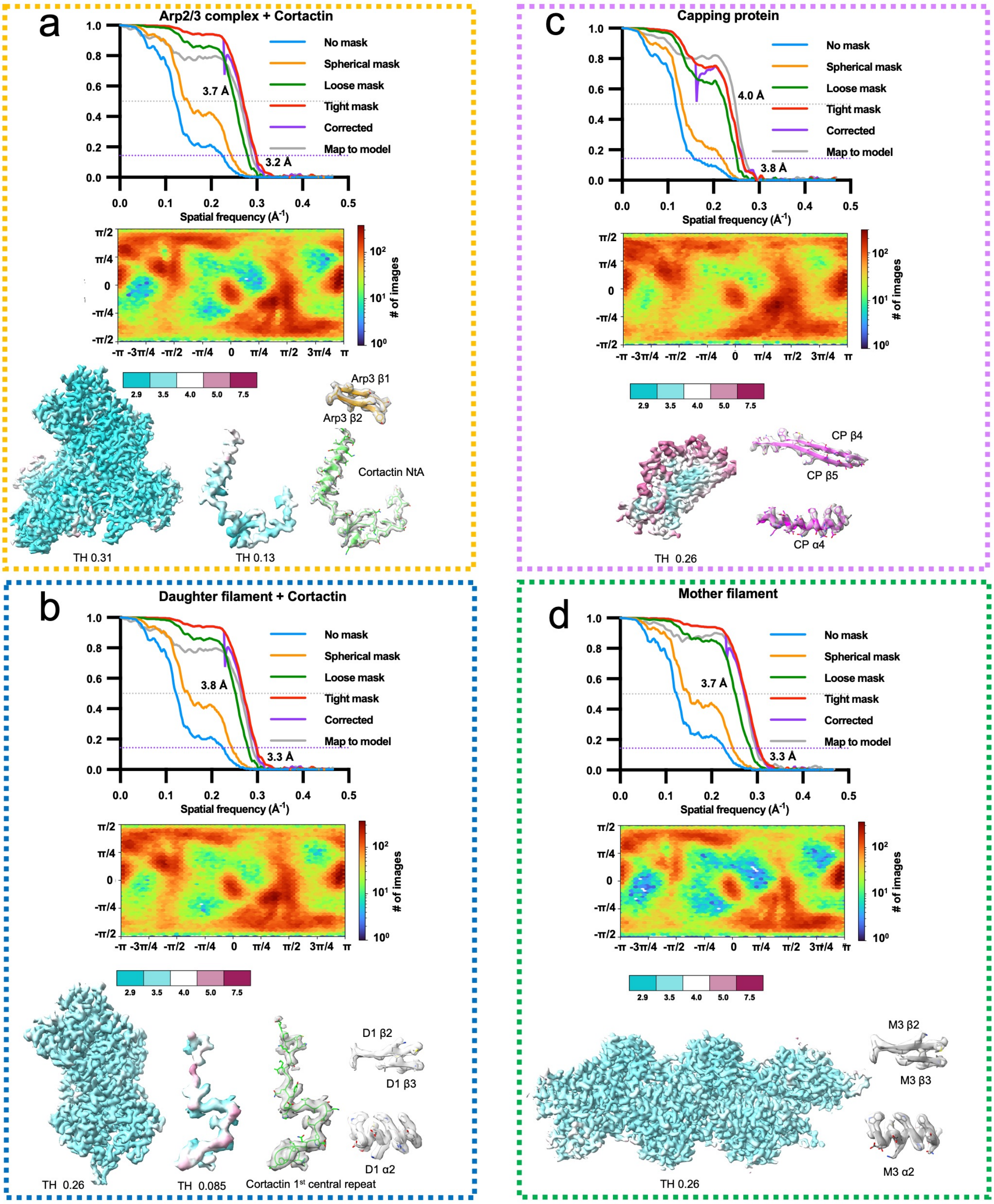


**Extended Data Fig 3. Cryo-EM data quality and validation of locally refined reconstructions.**

For each locally refined reconstruction shown in Extended Data Fig 2., half-map and map-model Fourier Shell Correlation (FSC), the angular distribution of particles used for 3D refinement, the well-resolved density used to generate the composite map coloured by local resolution and representative regions of the density map with the final model are shown. Thresholds (THs) are indicated. FSC Cut-off 0.143 was used for half-map resolution estimation. FSC Cut-off 0.5 was used for map-model resolution estimation and local resolution estimation. (a) Locally refined reconstruction of cortactin-bound Arp2/3 complex. (b) Locally refined reconstruction of daughter filament. (c) Locally refined reconstruction of capping protein. (d) Locally refined reconstruction of mother filament.

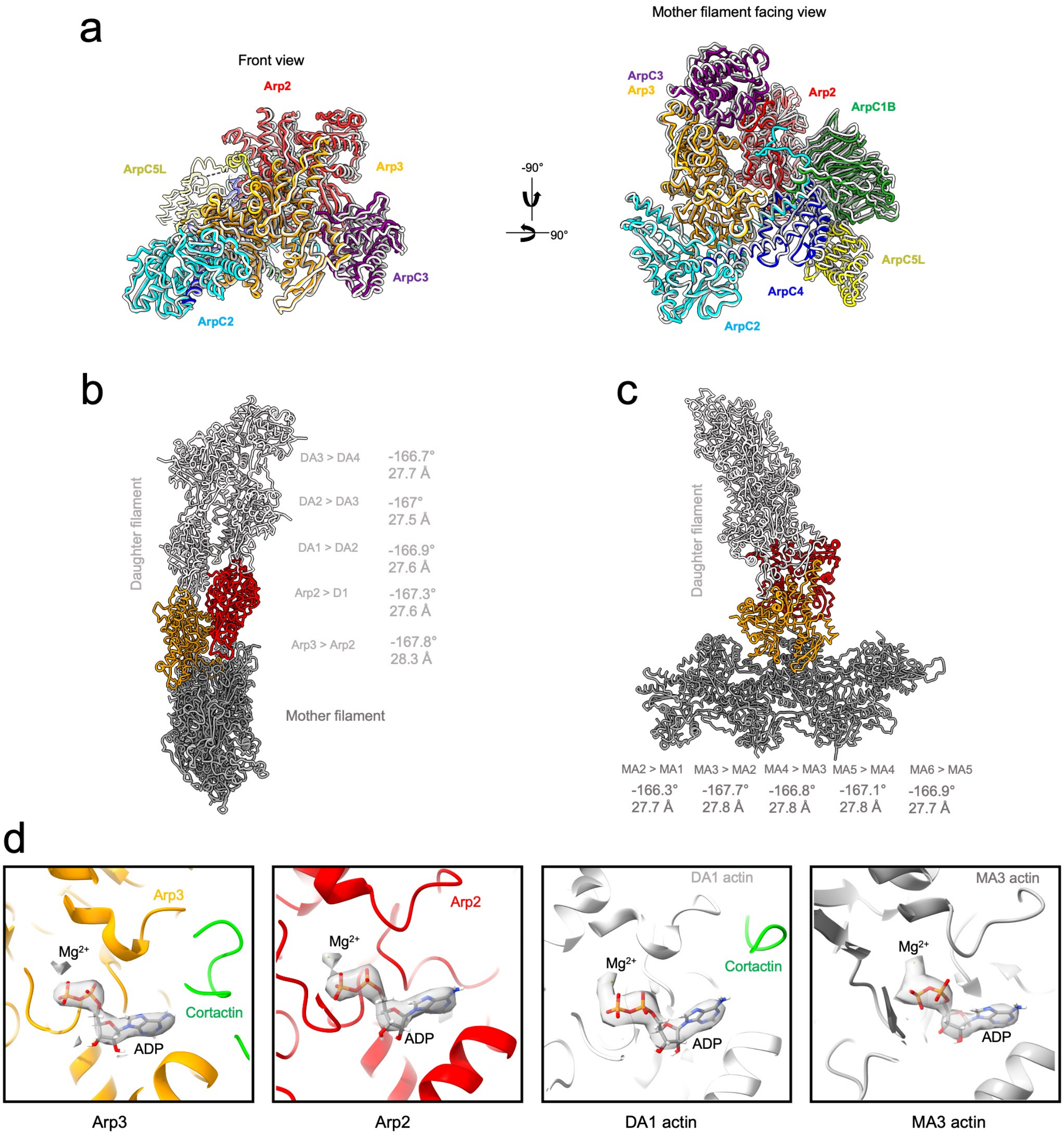


**Extended Data Fig 4. The activated Arp2/3 complex and ADP-F-actin in our cortactin-bound actin branch junction adopt canonical conformations.**

**a**) Structural alignment of cortactin-bound activated human Arp2/3 complex (this manuscript, subunits coloured) with activated bovine Arp2/3 complex (PDB 7tpt, subunits in light grey). Structures were aligned on ArpC2 (AA1-253) and Arp3 (AA1-36, AA60-153 and AA375-409). **b**) Arp2 and Arp3 in the activated Arp2/3 complex act as the template for daughter filament elongation. The canonical rise and twist between daughter filament actin subunits (DA1 – DA4) and/or Arp subunits are indicated. **c**) The canonical rise and twist between mother filament actin subunits (MA1 – MA6) are indicated and show no evidence of distortion within the mother filament upon branch formation. **d**) Density (transparent) and models of ADPs (in stick representation) and $Mg^{2+}$ (green dot) in Arp3, Arp2 and actin subunit DA1 in daughter and actin subunit MA3 in mother filament.

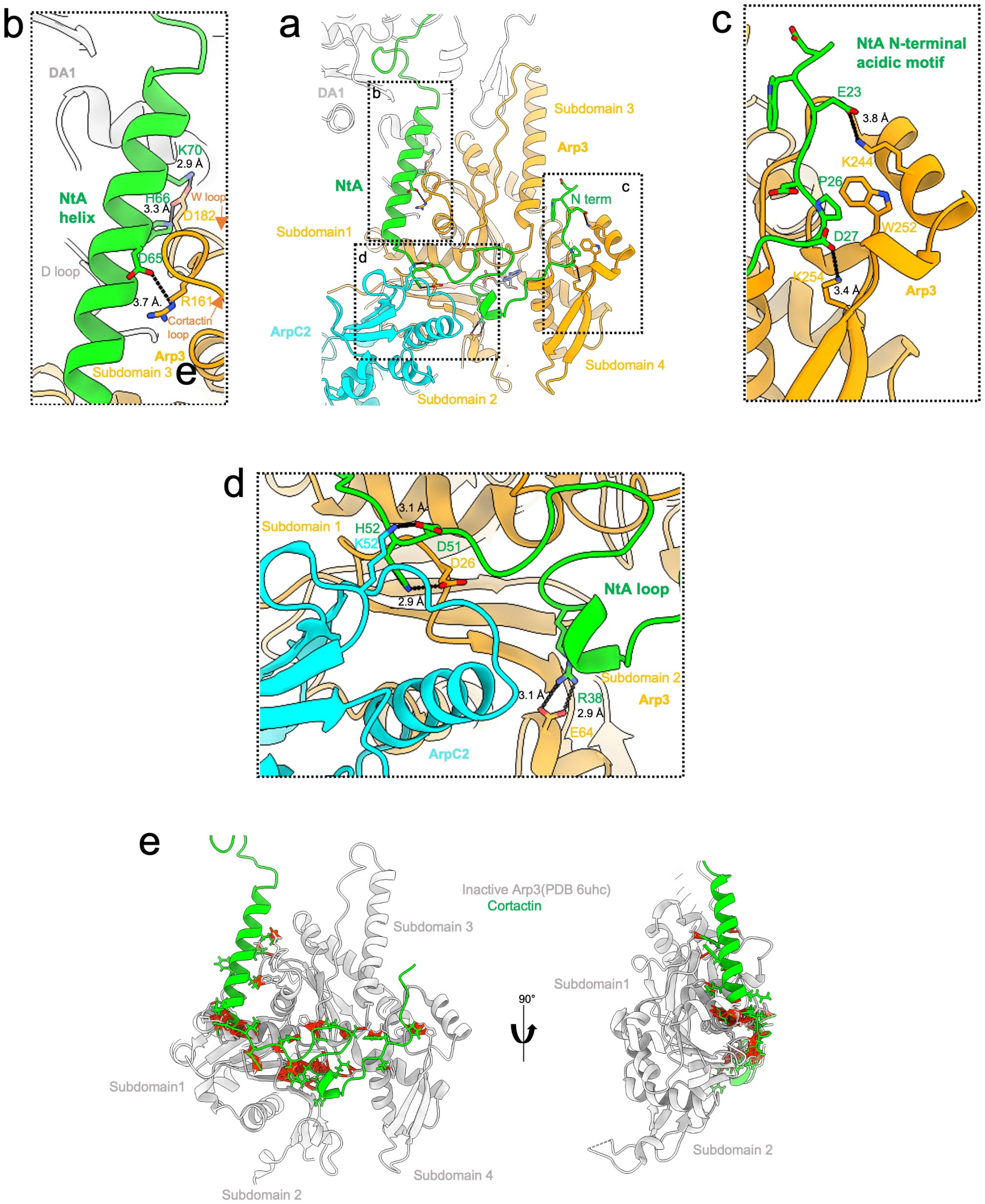


**Extended Data Fig 5. Interface details of the interactions between cortactin NtA and four subdomains of Arp3**

**a**) Overview of the cortactin NtA-Arp3-ARPC2 interface (similar to **Fig. 2a**). **b**) Details of the interaction between the cortactin NtA helix and Arp3 subdomain 3 (W loop and cortactin loop) of Arp3. **c**) Details of the interactions between NtA loop and subdomain 4. **d**) Details of the interaction between NtA loop and subdomain 1 and 2 of Arp3 and

ArpC2. **e**) Computational docking of our NtA structure onto the inactive Arp3 conformation generates structural clashes. Inactive Arp3 (from PDB 6uhc) is positioned by aligning on subdomains 3 and 4 of our activated Arp3 structure. Atom pairs with van der Waals overlap ≥ 0.7 Å (after subtracting 0.4 Å for H-bonding) were classified as clashes.

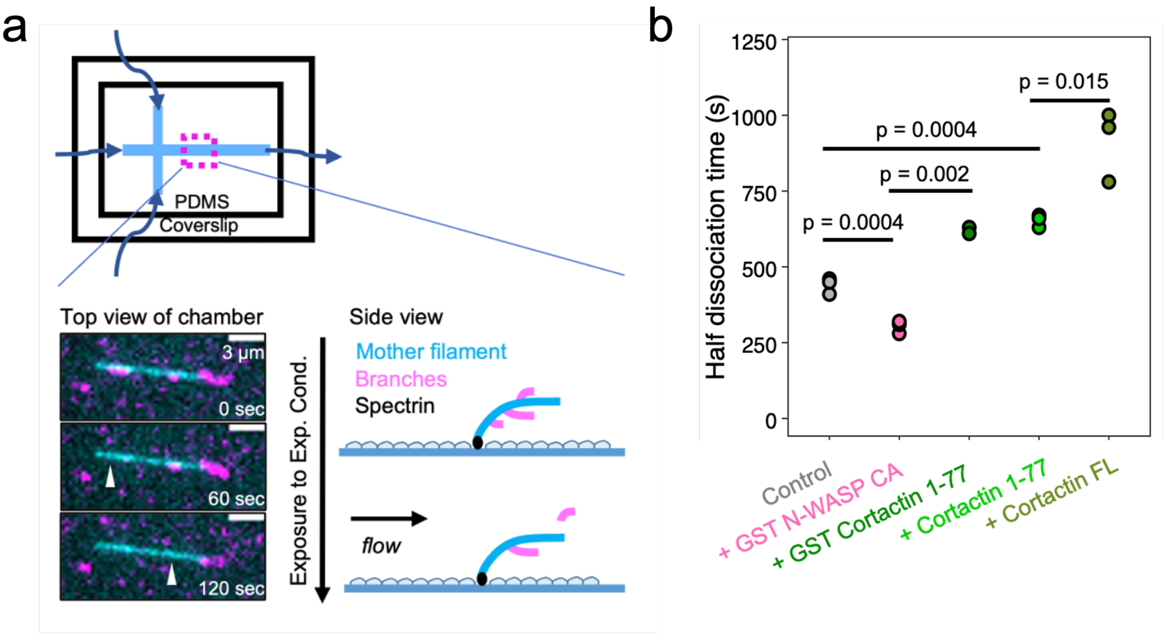


**Extended Data Fig 6. Microfluidics-based debranching assay shows the role of cortactin NtA in branch stabilization.**

**a**) A microfluidics setup was used to study the stability of Arp2/3-mediated branches. Actin filaments were attached to the surface of coverslip via their pointed ends. Actin branches were generated on top of the pre-existing filament using differently labelled actin. The dissociation of actin branches under the experimental conditions were observed and quantified. **b**) The time point when half of the actin branches have dissociated under different experimental conditions, taken from the same experiments as depicted in Fig. 4b. Each point represents the half dissociation time in an independent experiment. The p-value is estimated by unpaired t-test.

| | | | | |
|---|---|---|---|---|
| **Grid type** | C-flat 1.2/1.3 | | | |
| **Microscope** | Titan Krios | | | |
| **Detector and mode** | K3 super resolution mode (bin 2) | | | |
| **Collection software** | EPU | | | |
| **Magnification** | 81K | | | |
| **Voltage (kV)** | 300 | | | |
| **Electron exposure (e-/ Å$^2$)** | 49.4 | | | |
| **Exposure time (s)** | 3.8 | | | |
| **Dose rate (e-/pixel/s)** | 14.8 | | | |
| **Total frame** | 50 | | | |
| **Fraction dose (e-/ Å$^2$)** | 0.99 | | | |
| **Defocus range (µm)** | -0.9 to -2.4 | | | |
| **Pixel size (Å)** | 1.067 | | | |
| **Consensus reconstruction** | Mother filament consensus map (EMDB 17558) | Daughter filament consensus map (EMDB 17553) | | |
| **Particle number for final reconstruction** | 130,915 | 179,923 | | |
| **Map resolution (Å, FSC 0.143)** | 3.3 | 3.3 | | |
| **B factor** | -71.9 | -80.9 | | |
| **Local refined reconstruction** | Focus on mother filament (EMDB 17557) | Focus on Arp2/3 and cortactin (EMDB 17554) | Focus on daughter filament and cortactin (EMDB 17555) | Focus on capping protein (EMDB 17556) |
| **Particle number for final reconstruction** | 130,915 | 179,923 | 179,923 | 176,179 |
| **Map resolution (Å, FSC 0.143)** | 3.3 | 3.2 | 3.3 | 3.8 |
| **Local resolution range (Å)** | 3.0-6.5 | 2.9-5.2 | 3.0-7.3 | 3.5-6.9 |
| **B factor** | -75.6 | -78.4 | -81 | -112.3 |
| **Model resolution (Å, FSC 0.5)** | 3.7 | 3.7 | 3.8 | 4 |
| **Model composition** | PDB 8p94<br>Protein residue: 6380<br>Ligand: 12 MG, 12 ADP, 10 DTH, 10 EEP | | | |
| **B factors (Å$^2$)** | | | | |
| **Protein** | 76.4 | | | |
| **Ligand** | 33.3 | | | |
| **R.m.s. deviation** | | | | |
| **Bond lengths (Å)** | 0.007 | | | |
| **Bond angles (°)** | 1.491 | | | |
| **Molprobity score** | 1.65 | | | |
| **Clash score** | 9.6 | | | |
| **Rotamer outlier (%)** | 0 | | | |
| **Ramachandran Plot (%)** | | | | |

| | |
|---|---|
| **Favoured** | 97.2 |
| **Allowed** | 2.8 |
| **Disallowed** | 0 |

**Extended Data Table 1. Cryo-EM data collection, 3D image processing and model building statistics**
